# THE POTENTIAL ROLE OF TAURINE IN ACETAMINOPHEN HEPATOTOXIC INDUCED RATS

**DOI:** 10.1101/2024.08.23.609394

**Authors:** Isaac Olamide Babalola, Habeeb Kehinde Adekunle, Muhammed Damola Adigun

## Abstract

Taurine demonstrates an important cytoprotective function in the body, regulating oxidative and inflammation in various conditions. Taurine is actively synthesized in the hepatocyte, moreover, it confers protection to oxidative mediated injury in the liver. Acetaminophen, which has been shown to trigger oxidative related liver damage, is administered in its hepatotoxic dose to rats in this study. The preventive and regulatory role of taurine is investigated in this study. Twenty five adult male wistar rats were grouped into 5 distinct groups labeled the positive control group, negative control group, groups administered with taurine 12 hours prior acetaminophen administration, simultaneously with acetaminophen and an hour after acetaminophen was administered. The study indicated that taurine significantly reduced acetaminophen-induced liver injury in an in vivo rat model. Adding taurine either 12 hours before or at the time of acetaminophen treatment effectively prevented hepatocyte necrosis. Moreover, administering taurine 1 hour after the onset of acetaminophen-induced hepatotoxicity significantly improved liver injury, as evidenced by reduced hepatocyte necrosis, possibly through its unique cytoprotective properties such as antioxidant activity.

## Introduction

Taurine is a non-essential sulfur-containing amino acid found in high concentrations in various body cells, including hepatocytes, platelets, and leukocytes. Chemically, it is known as 2-aminoethanesulfonic acid (NH2CH2CH2SO3H), and primarily synthesized in the liver from the metabolism of methionine and cysteine [1]. Additionally, cells like neutrophils accumulate taurine through its uptake from the bloodstream. Taurine plays a crucial role in the immune system by acting as an antioxidant, protecting cells from oxidative stress. It is also very essential for cytoprotection and maintaining cellular homeostasis, particularly in response to acute and chronic inflammatory and oxidative stress conditions [2]. Oxidative stress, a major trigger of tissue damage associated with infections, inflammation, cancer, and aging, and primarily driven by reactive oxygen species (ROS) generated by activated leukocytes at sites of inflammation. Even though ROS are important for defending against pathogens, they can also contribute to tissue injury. In hepatic damage, severe oxidative stress could be from several factors including acetaminophen overdose [3,4]. Acetaminophen, commonly used as an analgesic and antipyretic for mild to moderate pain, can cause severe liver and kidney damage when overdosed. The toxicity begins with P450-mediated conversion of acetaminophen to NAPQI, which depletes glutathione and binds covalently to proteins and DNA in hepatic parenchymal cells, leading to liver injury. In addition to metabolic activation and NAPQI binding, reactive oxygen and nitrogen species, lipid peroxidation, mitochondrial dysfunction, disruption of calcium homeostasis, and necrosis also play roles in acetaminophen-induced hepatotoxicity [5,6,7]. This study aims to explore the potential protective effects of taurine against liver damage caused by acetaminophen. Identifying effective protective molecules is crucial for developing strategies to safeguard the liver during acute hepatic toxicity.

## Materials and Methods

### Study site

The study was conducted at the department of Medical Laboratory Science, Ladoke Akintola University of Technology Teaching Hospital, Ogbomoso, Oyo state, Nigeria.

### Study design

Twenty-five (25) adult male wistar rats (180 – 200g) were purchased from a breeding stock maintained in the animal house, College of Health Sciences, Ladoke Akintola University of Technology (LAUTECH), Ogbomosho, Oyo state. The animals were housed in plastic cages, and maintained at a temperature of 25 ± 5ºC. They were allowed to acclimatize to animal house conditions for one week before the commencement of research and were fed with a standard commercial feed pellet and provided with free access to clean water. The animals were allocated into five (5) groups, each of five (5) animals. All experimental procedures were conducted in accordance with National Institute of Health Guide for the Care and Use of Laboratory Animals (NIH, 1985) as well as Ethical Guidelines for the Use of Laboratory Animals in LAUTECH, Ogbomoso, Oyo state, Nigeria.

The wistar rats were randomly divided into five groups: a control group that received an equivalent volume of saline, an acetaminophen-only group, a group that received taurine 12 hours before acetaminophen injection, a group that received taurine and acetaminophen simultaneously, and a group that received taurine 1 h after acetaminophen injection therefore designated as; Group 1: (Negative control): 1 ml of Normal saline per kilogram per day for two weeks; Group 2 (Positive control): 1 ml of 500mg/kg/day of acetaminophen; Group 3: Normal rats administered 1ml of 200mg/kg taurine 12 h before 1 ml of 800mg/kg/day acetaminophen administration; Group 4: Normal rats administered 1ml of 200mg/kg taurine and 1ml 800 mg/kg acetaminophen simultaneously; Group 5: Normal rats administered 1ml of 200mg/kg taurine 1 h after 1 ml of 800mg/kg/day acetaminophen administration.

### Experimental protocol

30 mg/ml of acetaminophen was prepared in normal saline and orally administered in a hepatotoxic dose of 800 mg/kg. Taurine dissolved in normal saline (20 mg/ml) was also administered orally into the animals at a dose of 200mg/kg.

### Histological examination

After fourteen (14) days, precisely twenty-four (24) hours after treatment administration, the rats were sacrificed using cervical dislocation. The liver was removed and fixed in 10% formol saline for forty eight (48) hours to ensure sufficient fixation and prevent post mortem changes. Specimens were grossed and processed using the Automatic Tissue Processor (ATP) machine. The specimens were embedded with paraffin wax and sectioned using a rotary microtome. Sections were dewaxed in xylene and hydrated through 100%, 90% and 70% alcohol to water. They were subsequently stained in Harris haematoxylin for 15 minutes, held in running water for 2 minutes, differentiated in 1% acid alcohol, blued in running tap water for 10 minutes, counterstained in 1% eosin for 1 minute, dehydrated in ascending grades of alcohol, cleared in xylene and finally mounted with DPX. Ten satisfactory slides were viewed with a light microscope with two of the slides representing liver sections from each group.

## Results

**Table 1:** The histological report of liver sections from each experimental group.

| **ANIMAL GROUP** | **HISTOLOGICAL EXAMINATION** | **REMARK** |
| --- | --- | --- |
| **GROUP 1** | Sections show Liver tissue with preserved hepatocytes, portal triad and blood vessels. No atypical cell is seen. | Normal |
| **GROUP 2** | Section shows liver tissue with massive hepatic necrosis and congestion of sinusoid. There is severe interstitial lymphocytic infiltration. Also seen are distorted hepatocytes, widespread oedema and steatosis. | Severe necrosis |
| **GROUP 3** | Section shows liver tissue with mild lymphocytic infiltration of the interstitium. The hepatic architecture is moderately preserved with mild oedema. No necrosis or atypical cells seen. | Mild inflammation |
| **GROUP 4** | Section shows liver tissue with moderate to mild lymphocytic infiltration and mild centrizonal necrosis. | Moderate necroinflammation |
| **GROUP 5** | Section shows liver tissue with moderately preserved hepatic triad and blood vessels. The hepatocytes are moderately organized. Mild oedema seen. | Mild Oedema |

### Group 1: Negative control

**Figure 1:**
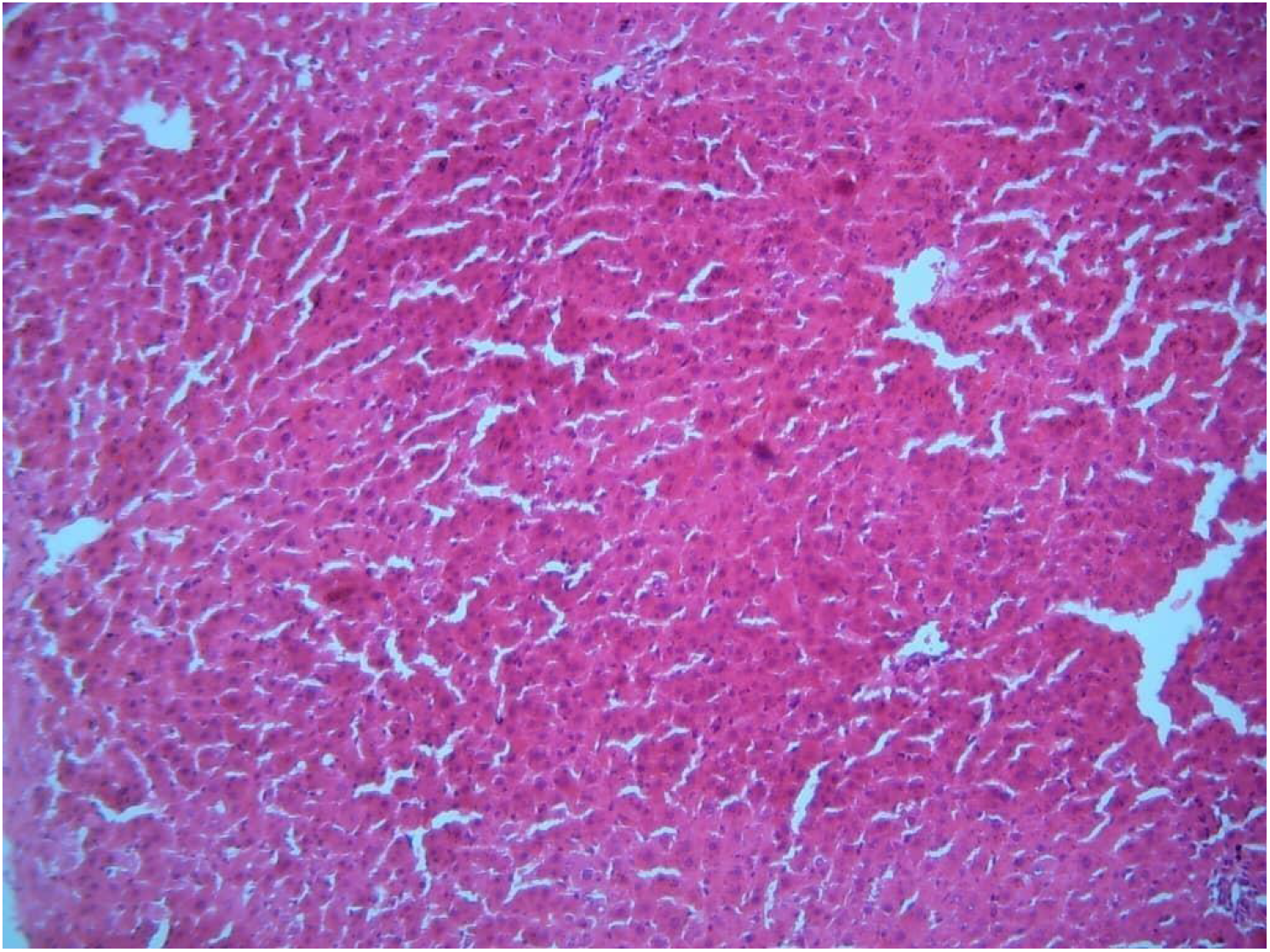
Liver Section for Negative Control (Mag-x100: Haematoxylin and Eosin) Sections show Liver tissue with preserved hepatocytes, portal triad and blood vessels. No atypical cell is seen

### Group 2 (Positive Control)

**Figure 2:**
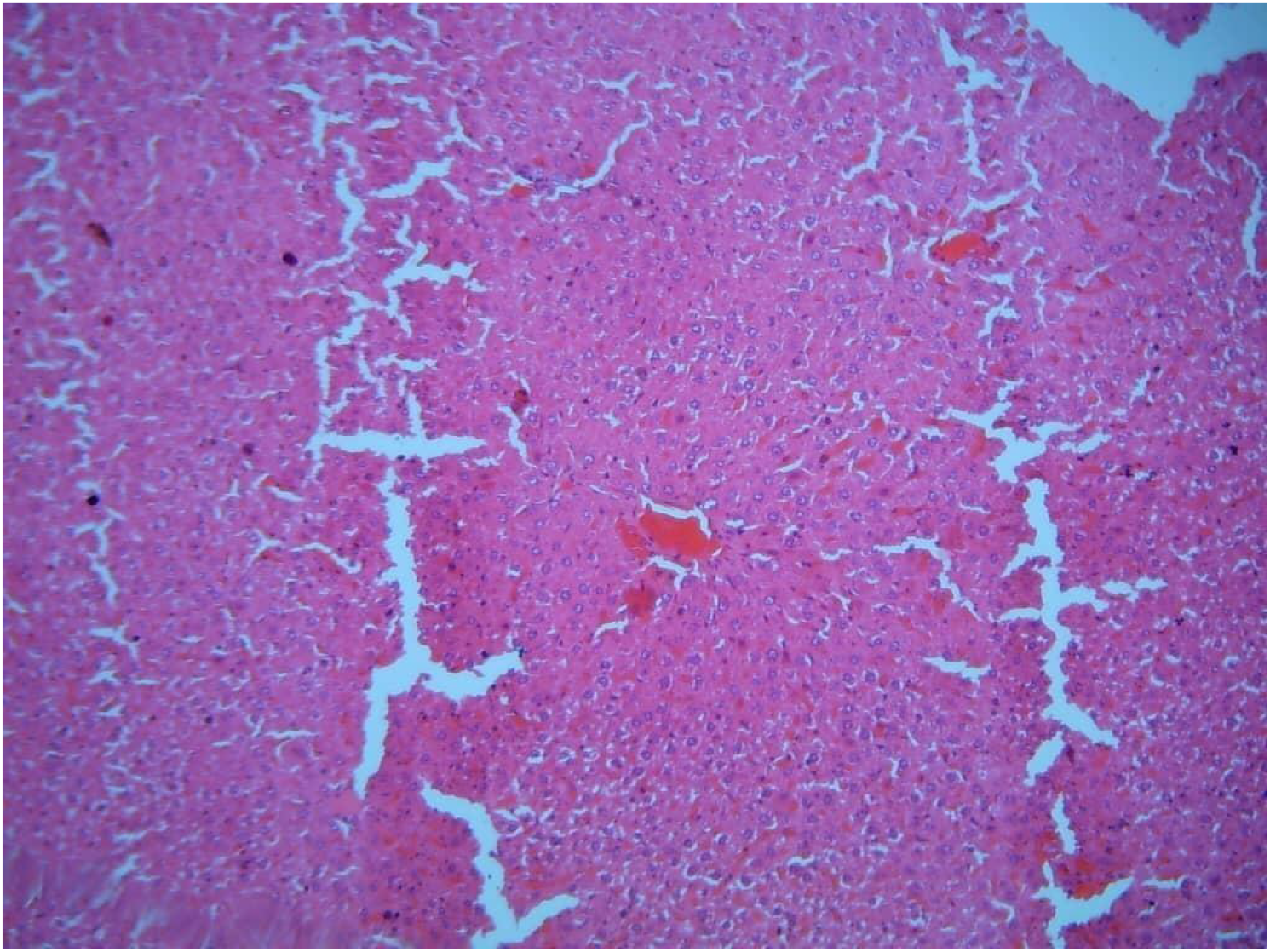
Liver Section for Positive Control (Mag-x100: Haematoxylin and Eosin) Section shows liver tissue with massive hepatic necrosis and congestion of sinusoid. There is severe interstitial lymphocytic infiltration. Also seen are distorted hepatocytes, widespread oedema and steatosis.

### Group 3 (Test 1)

**Figure 3:**
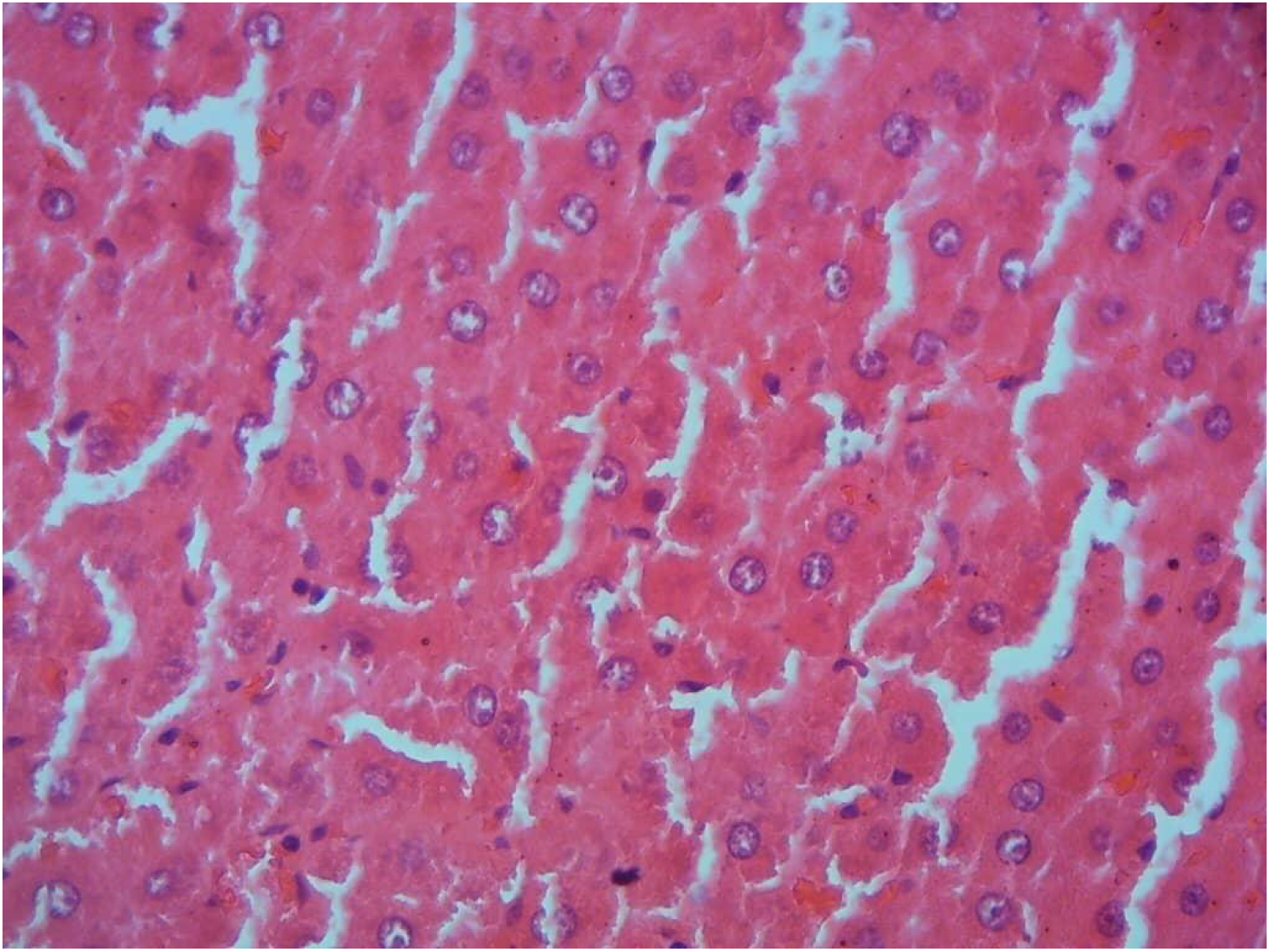
Liver Section for Test 1 (Mag-x100: Haematoxylin and Eosin) Section shows liver tissue with mild lymphocytic infiltration of the interstitium. The hepatic architecture is moderately preserved with mild oedema. No necrosis or atypical cells seen

### Group 4 (Test 2)

**Figure 4:**
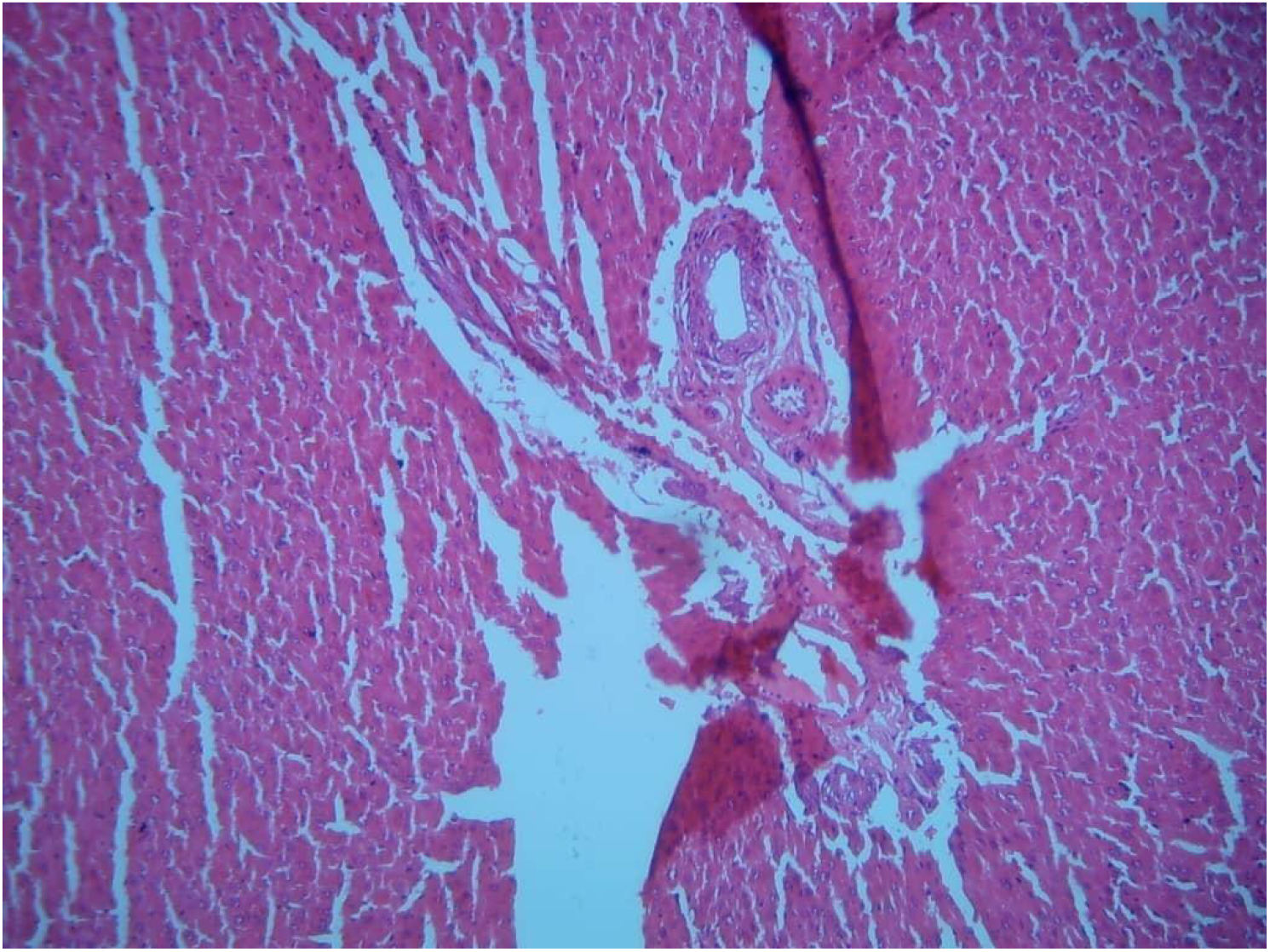
Liver Section for Test 2 (Mag-x100: Haematoxylin and Eosin) Section shows liver tissue with moderate to mild lymphocytic infiltration and mild centrizonal necrosis.

### Group 5 (Test 3)

**Figure 5:**
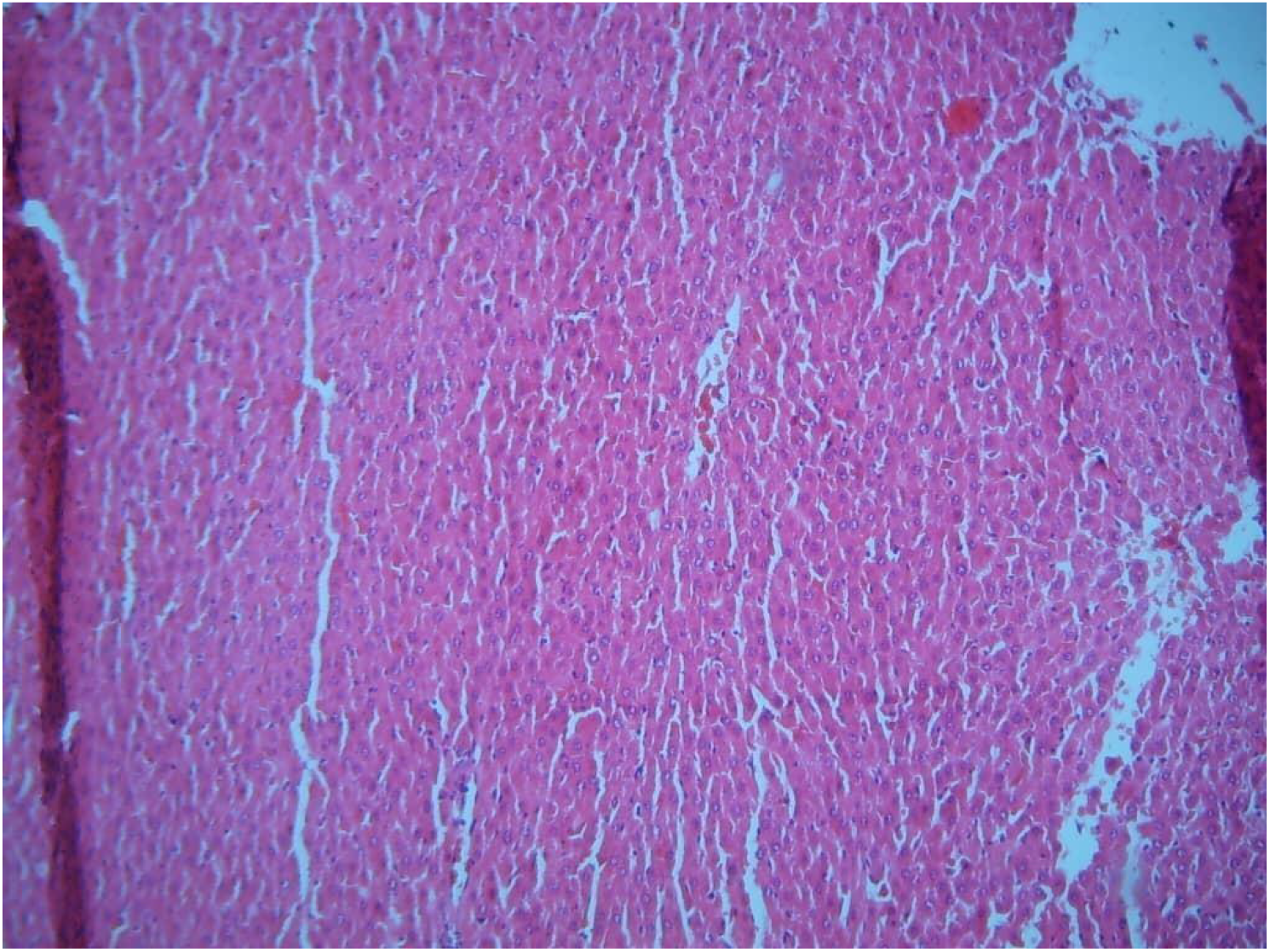
Liver Section for Test 3 (Mag-x100: Haematoxylin and Eosin) Section shows liver tissue with moderately preserved hepatic triad and blood vessels. The hepatocytes are moderately organized. Mild oedema seen.

## Discussion

Hepatocyte inflammation is a key process in acetaminophen-induced liver injury and typically precedes the onset of necrosis [9,10]. Liver damage from acetaminophen overdose may result from several mechanisms, including the generation of reactive oxygen species [11]. Studies have shown that taurine is important in cellular functions, such as membrane protection, antioxidant defense, and regulation of calcium levels [6,12]. In this study, administering a pharmacological dose of taurine significantly reduced acetaminophen-induced liver injury in an in vivo rat model. Adding taurine either 12 hours before or at the time of acetaminophen treatment effectively prevented hepatocyte necrosis. Moreover, administering taurine 1 hour after the onset of acetaminophen-induced hepatotoxicity significantly improved liver injury, as evidenced by reduced hepatocyte necrosis. These findings suggest that taurine has both prophylactic and therapeutic effects against acetaminophen-induced liver damage. Taurine has been used safely in humans at doses up to 5 grams [13], highlighting its potential as a preventive treatment for acetaminophen poisoning. Taurine administration, whether given before, simultaneously with, or after acetaminophen, significantly reduces hepatic injury by lowering hepatocellular enzyme release, decreasing hepatocyte apoptosis and necrosis, and reducing hepatic lipid peroxidation [14]. These findings suggest that taurine may play a crucial protective role in acetaminophen-induced liver injury, offering potential therapeutic benefits for mitigating its harmful effects. Additionally, taurine acts as a direct antioxidant by scavenging oxygen free radicals and inhibiting lipid peroxidation, as well as an indirect antioxidant by preventing increased membrane permeability due to oxidative damage in various tissues, including the liver [15]. Thus, taurine’s protective mechanism against acetaminophen-induced hepatotoxicity may be largely attributed to its antioxidant properties. Although this study did not directly demonstrate this effect, previous research has indicated that taurine significantly reduces acetaminophen-induced hepatic lipid peroxidation, which results from the interaction of reactive oxygen species with cell membrane phospholipids. This suggests that taurine helps mitigate acetaminophen-related oxidative damage in the liver. Similarly, other studies have highlighted taurine’s effectiveness in scavenging reactive oxygen intermediates and reducing lipid peroxidation, particularly in liver ischemia-reperfusion injury.

## Conclusion

This study demonstrate that administration of taurine has a prophylactic as well as a therapeutic role in preventing acetaminophen-induced hepatotoxicity, possibly through its unique cytoprotective properties such as antioxidant activity, inhibition of nitric oxide, and modulation of calcium homeostasis. These findings indicate that taurine deserves further consideration as a potential option for preventing acetaminophen overdose-related liver injury.

